# The Multifaceted Effects of Short-Term Acute Hypoxia Stress: Insights into the tolerance mechanism of *Propsilocerus akamusi* (Diptera: Chironomidae)

**DOI:** 10.1101/2023.06.05.543801

**Authors:** Yao Zhang, Qing-Ji Zhang, Wen-Bin Xu, Wei Zou, Xian-Ling Xiang, Zhi-Jun Gong, Yong-Jiu Cai

**Affiliations:** Key Laboratory of Watershed Geographic Sciences, Nanjing Institute of Geography and Limnology, Chinese Academy of Sciences, Nanjing 210008, China; School of Ecology and Environment, Anhui Normal University, Wuhu 241002, China; School of Geography and Ocean Science, Nanjing University, Nanjing 210023, China; Collaborative Innovation Center of Recovery and Reconstruction of Degraded Ecosystem in Wanjiang Basin Co-founded by Anhui Province and Ministry of Education, Wuhu 241002, P.R. China; College of Animal Sciences, Zhejiang University, Hangzhou, 310058, China

**Keywords:** Chironomidae, acute hypoxic stress, multi-omics analysis, energy metabolism, antioxidant mechanism

## Abstract

Plenty of freshwater species, especially macroinvertebrates that are essential to the provision of numerous ecosystem functions, encountered higher mortality due to acute hypoxia. However, within the family Chironomidae, a wide range of tolerance to hypoxia/anoxia is displayed and *Propsilocerus akamusi* depends on this great endurance to become a dominant species in eutrophic lakes. To further understand how *P. akamusi* responds to acute hypoxic stress, we used multi-omics analysis in combination with histomorphological characteristics and physiological indicators. To evaluate enzyme activity, the transcriptome and metabolome, and histomorphological characteristics, we set up two groups: a control group (DO 8.4mg/L) and a hypoxic group (DO 0.39mg/L). With blue-black chromatin, cell tightness, cell membrane invagination, and the production of apoptotic vesicles, tissue cells displayed typical apoptotic features. While lactate dehydrogenase (LDH), (Alcohol dehydrogenase) ADH, catalase (CAT), and Na+/K+ -ATPase (NKA) activities were dramatically enhanced under hypoxic stress, glycogen content, and superoxide dismutase (SOD) activities were significantly reduced compared to the control group. The above results were further supported by the joint analysis of the transcriptome and metabolome, which further revealed that in addition to carbohydrates, including glycogen, the energy metabolism of the fatty acid, trehalose, and glyoxylate cycles is also included. Furthermore, we also revealed the ethanol tested in hypoxic stress should derive from symbiodinium of *P akamusi.* Understanding the processes which enable *P. akamusi* to survive lengthy periods of hypoxia in eutrophic lakes might help to design sensitive biomonitoring procedures, and this species has the potential to be used as an effective eutrophication indicator.

## 1. Introduction

Hypoxic zones have been rapidly expanding in both space and frequency in freshwater ecosystems, as a result of rising nutrient loadings and global warming(Conley *et al*., 2009; Diaz, 2001; Jenny *et al*., 2014; Jenny *et al*., 2016; Rabalais *et al*., 2010; Wang *et al*., 2016; Zhang *et al*., 2022). However, acute hypoxia leads to higher mortality rates in numerous freshwater species, especially macroinvertebrates, which are crucial to the provision of many ecosystem functions(Galic *et al*., 2019; Pardo *et al*., 2016). The makeup of the macroinvertebrate community has also significantly deteriorated as hypoxia changed the metabolic response (metabolomes) and maybe other biochemical reactions, such as the status of oxidative stress (Saari *et al*., 2018; Verberk *et al*., 2013).

Hypoxia stress not only led to significant tissue damage but also induced cell apoptosis and the expression of hypoxia-inducing genes and proteins has been identified in macroinvertebrates, focusing on marine organisms (Andreyeva *et al*., 2021; Felix-Portillo *et al*., 2016; Geihs *et al*., 2014; Grimes *et al*., 2020; Hu *et al*., 2022; Sun *et al*., 2019; Wang *et al*., 2022; Wang *et al*., 2019; Wang *et al*., 2019). Studies have also shown in scallops, crabs, and clams that energy-consuming processes (including protein turnover and ion transport) are suppressed by hypoxia stress, which is indicated by a dramatic decrease in the activity of Na+/K+ -ATPase NKA (Ivanina et al., 2016; Lucu, 2020). However, it is unclear the effects of hypoxia on freshwater macroinvertebrates, especially the family Chironomidae with a wide range of tolerance to hypoxia (Frank, 1983; Hamburger *et al*., 1994; Sun *et al*., 2018). In addition, hypoxia stress also altered the metabolite content of insect, relating to energy and antioxidants(Campbell *et al*., 2019; Cui *et al*., 2017). Hypoxic stress increases reactive oxygen species (ROS) production in the body by disrupting the electron transport chain and the antioxidant defense system, including enzymes such as superoxide dismutase (SOD) and catalase (CAT), is activated to avoid ROS harmful effects on biomolecules(Correia et al., 2003; Via et al., 1998). For instance, In the aquatic insect *Belostoma elegans*, hypoxic conditions have been shown by Lavaras et al. (2017) to be one of the factors affecting antioxidant enzyme activity (Lavarías et al., 2017). The evidence indicates that ethanol is the only byproduct of glycogen degradation from *Chironomus thummi* and *Culex pipiens* in hypoxic stress, and it is catalyzed by the enzyme alcohol dehydrogenase (ADH) or diffused into the surrounding water (Redecker et al., 1988; Zebe, 1991). However, the study on the *Chaoborus crystallinus* found that large quantities of alanine are accumulated, while lactate is produced as a minor end product under anoxic conditions (Englisch et al., 1982).

Chironomids are widely dispersed, simple to cultivate, have brief life cycles, and are also adapted to a variety of environmental rigors, including desiccation, anoxia, high temperature, freezing, eutrophication, and chemical pollution (Choi, 2004). Thus, as bioindicators, they are a suitable taxon to explore the adaptation mechanisms required to endure environmental stress (Lencioni *et al*., 2008; Rosenberg, 1992). Chironomid midge larvae possess a wide range of tolerance to hypoxia, due to extracellular hemoglobins (Hbs) in monomeric and dimeric forms floating in their hemolymph (Hamburger *et al*., 1994; Nath, 2018). Especially, *P. akamusi* of Chironomidae frequently predominates macroinvertebrates in many eutrophic lakes and demonstrated incredibly low fuel consumption at a stage of estivation when the larvae lived in deep sediment and engaged in anaerobic respiration, with ethanol as a major metabolite, to withstand prolonged hypoxia (Gong *et al*., 2008; Hirabayashi *et al*., 2004; Zou *et al*., 2019). However, the mechanisms of how *P. akamusi* survives for a prolonged period of time in severe hypoxic environment are still unclear.

Approaches to *P. akamusi* have mostly focused on the transcriptome, proteome, or metabolome(Liu et al., 2022; Liu et al., 2022; Sun et al., 2022; Yan et al., 2020; Zheng et al., 2017). However, single-omics cannot systematically explain the rapid biochemical reactions and metabolic changes that are expected for acute hypoxia(Hasin *et al*., 2017; Manzoni *et al*., 2018). For this reason, we conducted a comprehensive comparative metabolome–transcriptome analysis to understand how P. akamusi, dominant macroinvertebrate species in many eutrophic lakes, endures prolonged hypoxia(Zou et al., 2018). Thus, we combine physiological indicators and histomorphology observations with metabolome–transcriptome analysis to comprehensively assess histomorphological features, energy metabolism mechanism, and antioxidant mechanism of *P. akamusi* larvae exposed to acute hypoxia.

## 2. Materials and methods

### 2.1 Experimental animals, hypoxia challenge, and tissue sample preparation

The larvae of *P. akamusi* were collected in the Taihu Lake basin, and the viable P. akamusi larvae were put in culture boxes for further studies. P. akamusi larvae were put in 12 culture boxes, and a hypoxic experimental group and a hypoxic group with three biological replicates each were established. The temperature was kept constant at 22-25°C throughout the experiment, and the oxygen concentration in the control group was not less than 8.0mg/L. The hypoxic method used was nitrogen-filled, and the oxygen content was checked at all times to ensure that it did not exceed 0.5mg/L. The alcohol content in the culture fluid was also checked, and once ethanol was detected, samples were immediately taken and placed in liquid nitrogen bottles. At this point, the time was set to 0h, after which samples were taken at 3h, 6h, 12h, 24h, 48h, 72h, and 96h. At 0 h, 12 h, 24 h, 48 h, 72 h, and 96 h after hypoxic stress, 15 whole *P. akamusi* larvae were sampled from each parallel bucket at each time point. The larvae were put into 1.5 mL EP tubes, briefly submerged in liquid nitrogen for short-term storage, and then kept at -80°C in an ultra-low temperature refrigerator.

### 2.2 Transcriptome analysis

Utilizing the TRIzol reagent, which is available from Invitrogen in California, USA, total RNA was isolated from P. akamusi larvae. Total RNA quantity and integrity were assessed using the Agilent Bioanalyzer 2100 equipment (Agilent, CA, USA). Each sample was chosen to have 1.5 g of high-quality RNA for the investigation. At a high temperature, divalent cations break the mRNA into minute pieces. The mRNA Seq preparation sample kit (Illumina, San Diego, USA) instructions were then followed to construct the final cDNA library utilizing reverse transcription. On the Illumina NovaseqTM 6000 platform (LC Sciences, USA), paired-end DNA sequencing was carried out according to the vendor’s suggested technique. Clean data (clean reads) were chosen from the raw data (raw reads) to explore the patterns of gene expression in P. akamusi larvae under hypoxic stress, then the sequence quality was checked using FastQC (https://fastqc.com/). With Trinity 2.4.0, clean data were then put together into a transcriptome for reference (Grabherr *et al*., 2011). The “gene” sequence (also known as the Unigene) was chosen as the cluster’s longest transcript. The Basic Local Alignment Search Tool (BLAST) was used for transcriptome annotation, while the DIAMOND technique searches numerous databases with an E-value of 1×e^-5^. We employed Blast2GO with NR annotation for Gene Ontology (GO) annotation and the default settings of KASS for KEGG pathway analysis. The comparison of the hypoxic and control groups was done using an evaluation of differential gene expression. To measure the amount of Unigene expression, salmon was used (Mortazavi *et al*., 2008; Patro *et al*., 2017). The R package edgeR (Robinson et al., 2010) was used to identify the differentially expressed unigenes with statistically significant (P value < 0.05). using Perl scripts in R language, the DEGs were assessed for GO enrichment and KEGG pathway enrichment based on hypergeometric distributions. The raw data can be viewed in the SRA (Sequence Read Archive) database, number: PRJNA972485.

### 2.3 Metabolome analysis

To get baseline correction, maximum alignment, maximum detection, accurate masses, and normalized intensity of the peak, Agilent MSD Chemstation (version E.02.00.493) using the default settings was chosen(Smith *et al*., 2006). Then, maximum acquisition and deconvolution were carried out using the automated mass spectrometry deconvolution recognition system (AMDIS). The identification of metabolites was then completed by contrasting mass fragment patterns and retention times across several databases. Differentiating metabolites between the Q and W groups were mapped to their corresponding metabolic pathways with the KEGG pathway-based MetPA online tool (http://metpa.metabolomics.ca/). Only metabolic pathways with -lg (p) < 1.301 were preserved based on hypergeometric testing.

### 2.4 Tissue enzyme activity and glycogen assays

The kits for the determination of glycogen content, ADH activity, LDH activity, CAT activity, SOD activity, and NKA activity were purchased from Nanjing Jiancheng Biological Company, and the enzyme activities and glycogen content of the hypoxic and control groups were determined using tissues as experimental samples. The specific operation procedure and calculation formula were referred to in the instruction manual.

### 2.5 HE staining

The P. akamusi larvae were washed in pre-cooled saline at 4 °C, and the tissue was then chopped into small pieces and stored on ice before being placed in 10% formaldehyde solution and fixed for one day and one night before being placed in ethanol for three hours to dehydrate it. After being sectioned and paraffin-embedded, the slides were dried on a staining rack for 20 minutes before being submerged in a solution of xylene and ethanol for 12 minutes. The slides were then dehydrated in ethanol for six minutes, dehydrated in xylene for twenty minutes, washed in xylene, stained in hematoxylin for two minutes, rinsed under low-flow water for four minutes, and then soaked in eosin solution for twenty seconds. The slides were then sealed with gum, dried, and examined under a biological microscope. After drying, the films were examined under a biological microscope. (Ma *et al*., 2021).

### 2.6 Data analysis

To examine the significance of differences between groups, the data were analyzed using the statistical program SPSS 25.0 for one-way ANOVA, the chi-square test, and LSD or Duncan’s multiple comparisons depending on the findings of the chi-square test, respectively (p< 0.05 was considered significant). ORIGIN software was used to plot the results, which were presented as mean standard deviation (x ± SD).

## 3. Results

### 3.1 Effect of hypoxic stress on Histomorphological feature and NKA activity in *P. akamusi*

Compared with the control group (Fig. 1A), the *P. akamusi* staining in the hypoxic group was darker and tighter (Fig. 1C). The tissue cells in the hypoxic group were haphazardly distributed with different cell morphologies, significantly smaller in size than the control group, and displayed a tighter state (Fig. 1D). In contrast, tissue cells in the control group were neatly arranged, maintaining normal cell morphology and volume (Fig. 1B). The tissue cells in the hypoxic group tended to be rounded with blue-black chromatin, and some of the cell membranes were crinkled and invaginated, indicating apoptosis (Fig. 1F black arrows). Some tissues displayed apoptotic vesicles (Fig. 1G star-shaped marker cells). Under hypoxic stress, a significant number of P. *akamusi* tissues exhibited morphological characteristics of apoptosis.

**Fig. 1.**
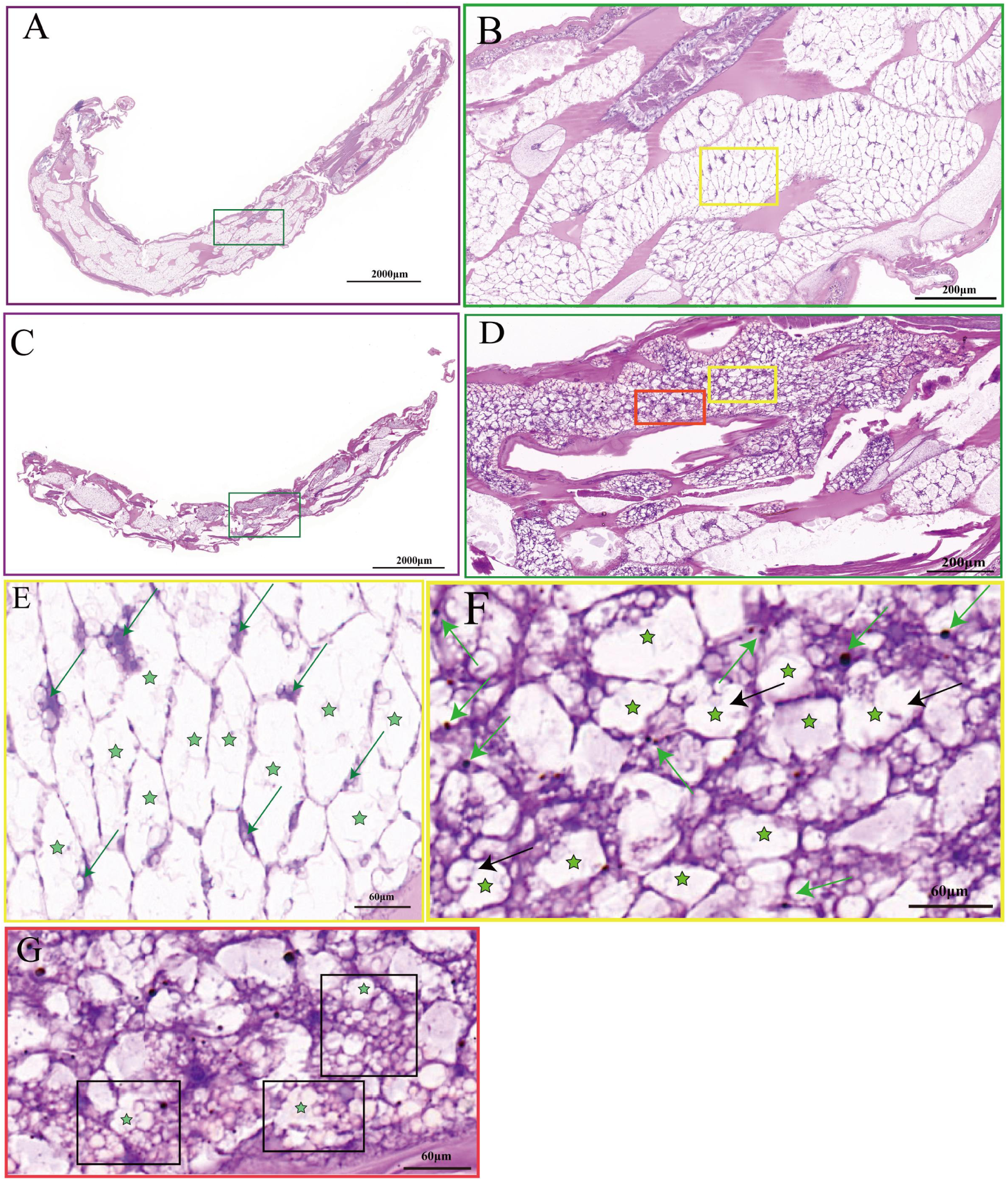
Histomorphological feature of the *P. akamusi* Note: Control group includes A (upper; 3×), B (upper; 30×), E (upper; 90×), Hypoxic group includes C (upper; 3×), D (upper; 30×), F (upper; 90×), G (upper; 90×).

The hypoxic and control groups showed similar trends and significant changes. During hypoxic stress, the NKA activity of the hypoxic group was significantly higher than that of the control group (Fig 2). The NKA activity of the control group displayed an overall trend of increasing and then fluctuating downward, with significant changes. Specifically, it increased significantly at 12h, decreased significantly, increased significantly again at 48h, and decreased significantly again at 96h, with activity similar to the initial value. In the hypoxic group, NKA activity showed a fluctuating upward trend followed by a downward trend, with significant changes. Specifically, it increased significantly at 12h, decreased significantly at 24h, then increased significantly and reached a maximum at 72h, and decreased significantly and reached a minimum at 96h, with activity similar to the initial value.

**Fig 2.**
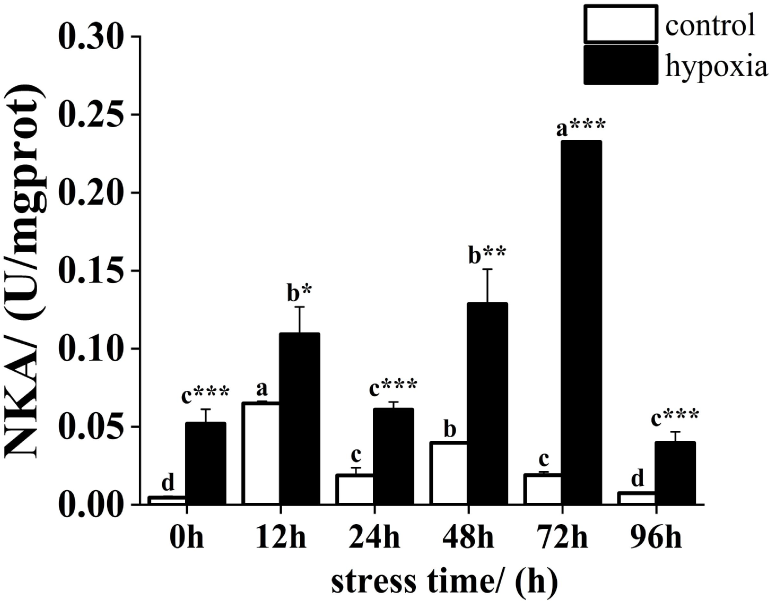
Changes in NKA activity of the *P. akamusi* under hypoxic stress Note: Different letter superscripts indicate significant differences within the control or hypoxic group in different stress time (p<0.05) * indicate significant difference between groups at the same time point, * indicates p<0.05, ** indicates p<0.01, *** indicates p<0.001, below same.

### 3.2 Effect of hypoxic stress on energy metabolism in the *P. akamusi*

The glycogen content in the control group was significantly higher than that in the hypoxic group during hypoxic stress. The glycogen content in the control group tended to decrease overall, with a significant decrease at 12h and 96h and reaching a minimum value at 96h, which was significantly lower than the initial value. In the hypoxic group, there was a decreasing trend followed by an increasing trend with a significant change. its content reached its lowest value at 48h and was significantly lower than the initial value at 96h (Fig 3. A). LDH activity was significantly higher in the hypoxic group than in the control group at all times of hypoxic stress, except for 12h and 48h. LDH activity in the control group decreased significantly at 24h, increased significantly at 48h, and reached a maximum value, then decreased significantly and reached a minimum value at 96h, and this value was significantly lower than the initial value. In the hypoxic group, LDH activity showed a fluctuating upward trend. its activity increased significantly at 24h, but decreased significantly at 72h, then increased significantly and reached a maximum at 96h, and the value was about 3.5 times higher than the initial value (Fig 3 B). Except for 96h, the ADH activity of the hypoxic group was significantly higher than the control group at different stress times. overall, the ADH activity of the control group showed fluctuating but non-significant changes. the ADH activity of the hypoxic group showed a fluctuating decreasing trend. during 48h of hypoxic stress, the ADH activity continued to decrease, but at 72h the ADH activity increased significantly, then decreased significantly and reached the minimum value at 96h, and the value was significantly lower than the initial value (Fig 3 C).

**Fig 3.**
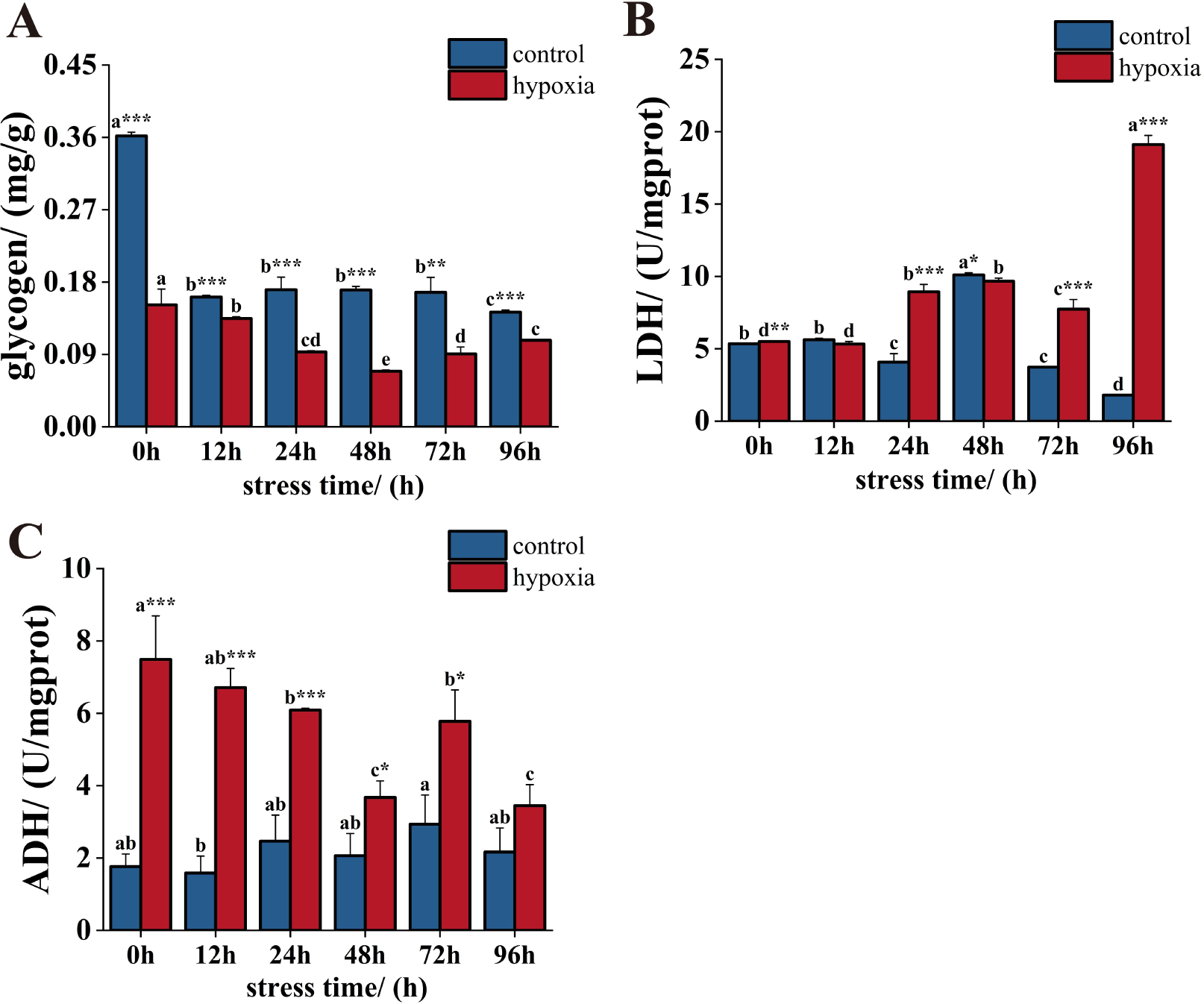
Changes in glycogen content(A), ADH activity(B) and LDH activity (C) of the *P. akamusi* under hypoxic stress

### 3.3 Effect of hypoxic stress on antioxidant enzyme activity in *P. akamusi*

During hypoxic stress, the SOD activity of the control group was significantly higher than that of the hypoxic group except for 0h (Fig 4A). The SOD activity of the control group displayed an overall fluctuating trend with significant change. Specifically, it increased significantly at 12h, decreased significantly, and reached a low value at 48h, then increased significantly and reached a maximum value at 96h, which was significantly higher than the initial value. In contrast, the SOD activity of the hypoxic group showed a trend of increasing and then decreasing, with a significant decrease at 24h and reaching its lowest value, followed by a significant increase at 72h and stabilization. However, at 96h, the SOD activity was significantly lower than the initial value. The CAT activity of the hypoxic group was significantly higher than that of the control group at different stress times (Fig 4 B). In the control group, CAT activity increased significantly at 12h and reached a maximum value, then decreased significantly and reached a minimum value at 48h, but increased significantly at 72h and then stabilized, and CAT activity was similar to the initial value at 96h.

**Fig 4.**
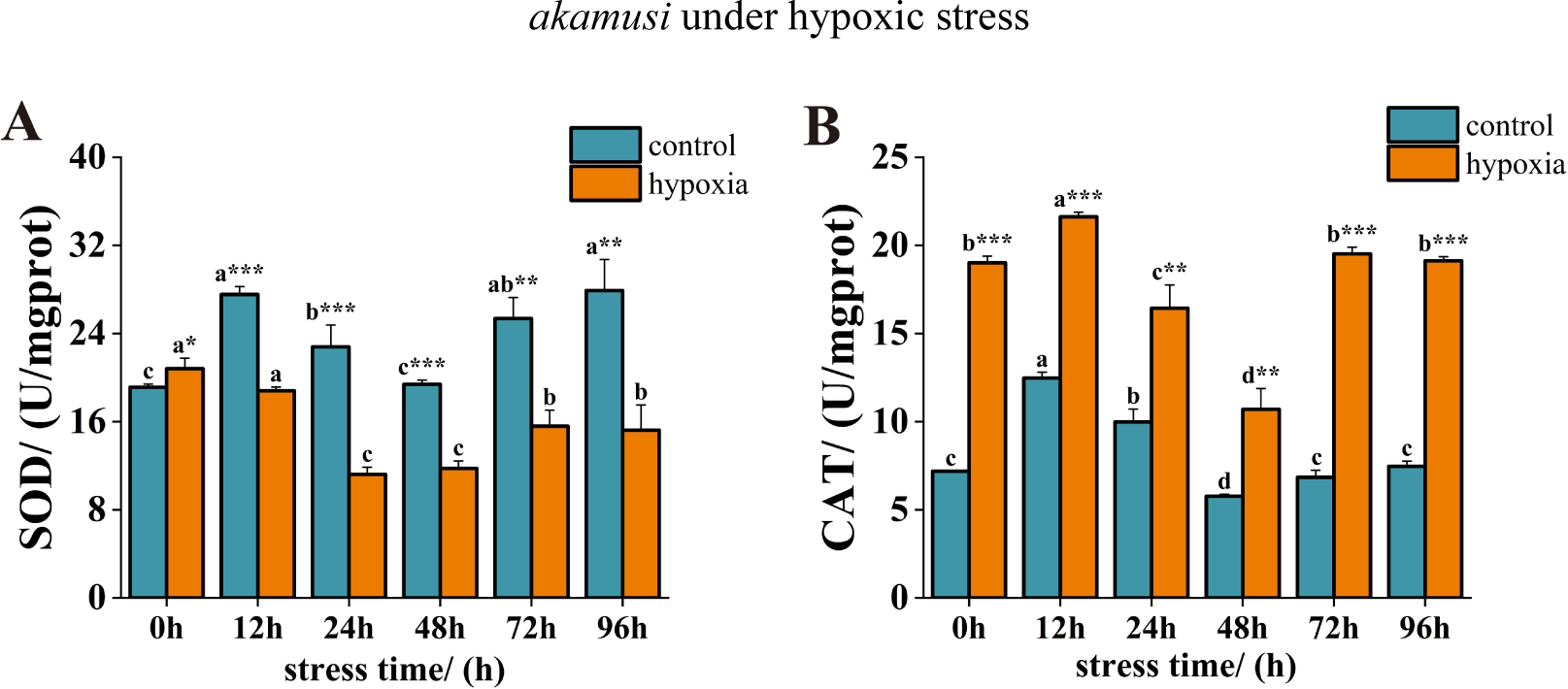
Changes in SOD activity(A) and CAT activity(B) of the *P. akamusi* under hypoxic stress

### 3.4 GO enrichment and KEGG enrichment analysis of all differentially expressed genes

GO annotation enrichment was performed to analyze differentially expressed genes in *P. akamusi* under hypoxic stress (Fig 5). The results revealed significant enrichment of metabolic and catabolic processes involved in differentiation, lipid metabolic process, carbohydrate derivative metabolic progress, and trehalose metabolic progress from a biological process ontology perspective. In terms of molecular function perspective, catalytic activity, oxidoreductase activity, transferase activity, transmembrane receptor protein tyrosine kinase activity, carbohydrate kinase activity, and trehalas activity were observed. For cellular components, membrane-related components, such as vesicle membrane, endomembrane system, and extracellular region were significantly enriched.

**Fig 5.**
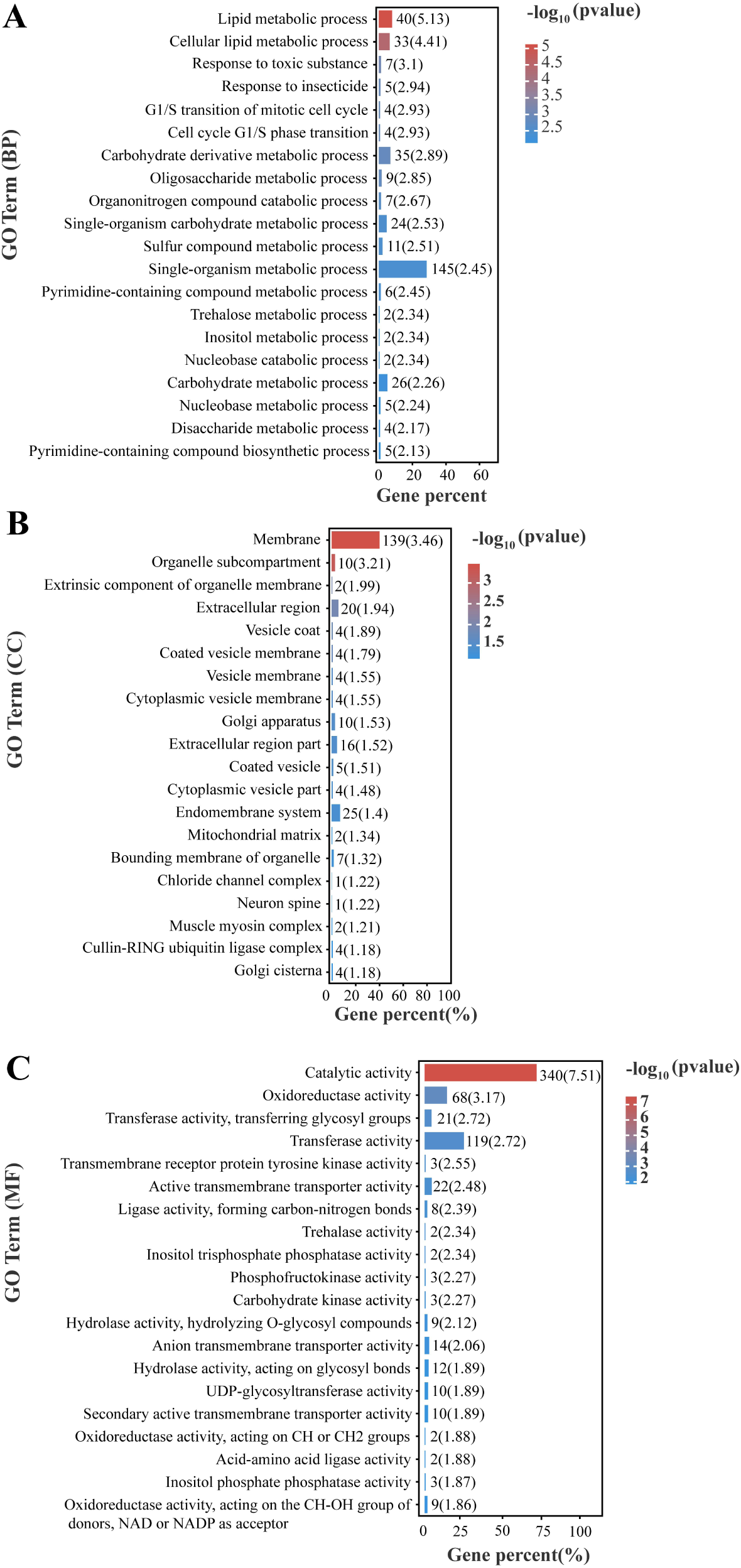
Top 20 GO terms enrichment (biological processes, cellular component and molecular function) of all DEGs under hypoxia stress.

KEGG annotation enrichment was used to analyze the differentially expressed genes in *P. akamusi* under hypoxia stress (Fig 6). Several pathways were significantly altered during hypoxia stress, including metabolic pathways, drug metabolism pathways, pyrimidine metabolism, porphyrin and chlorophyll metabolism pathways, ABC transporters, and cytochrome p450-related pathways. The differentially expressed genes in these pathways include ATP-binding cassette, subfamily C (CFTR/MRP), member 4(ABCC4) ATP-binding cassette, subfamily G (WHITE), member 1(ABCG1), ATP-binding cassette, subfamily G (WHITE), member 4(ABCG4), dimethylaniline monooxygenase, (N-oxide forming)/ hypotaurine monooxygenase (FMO), glucuronosyl transferase (UGT), etc.

**Fig 6.**
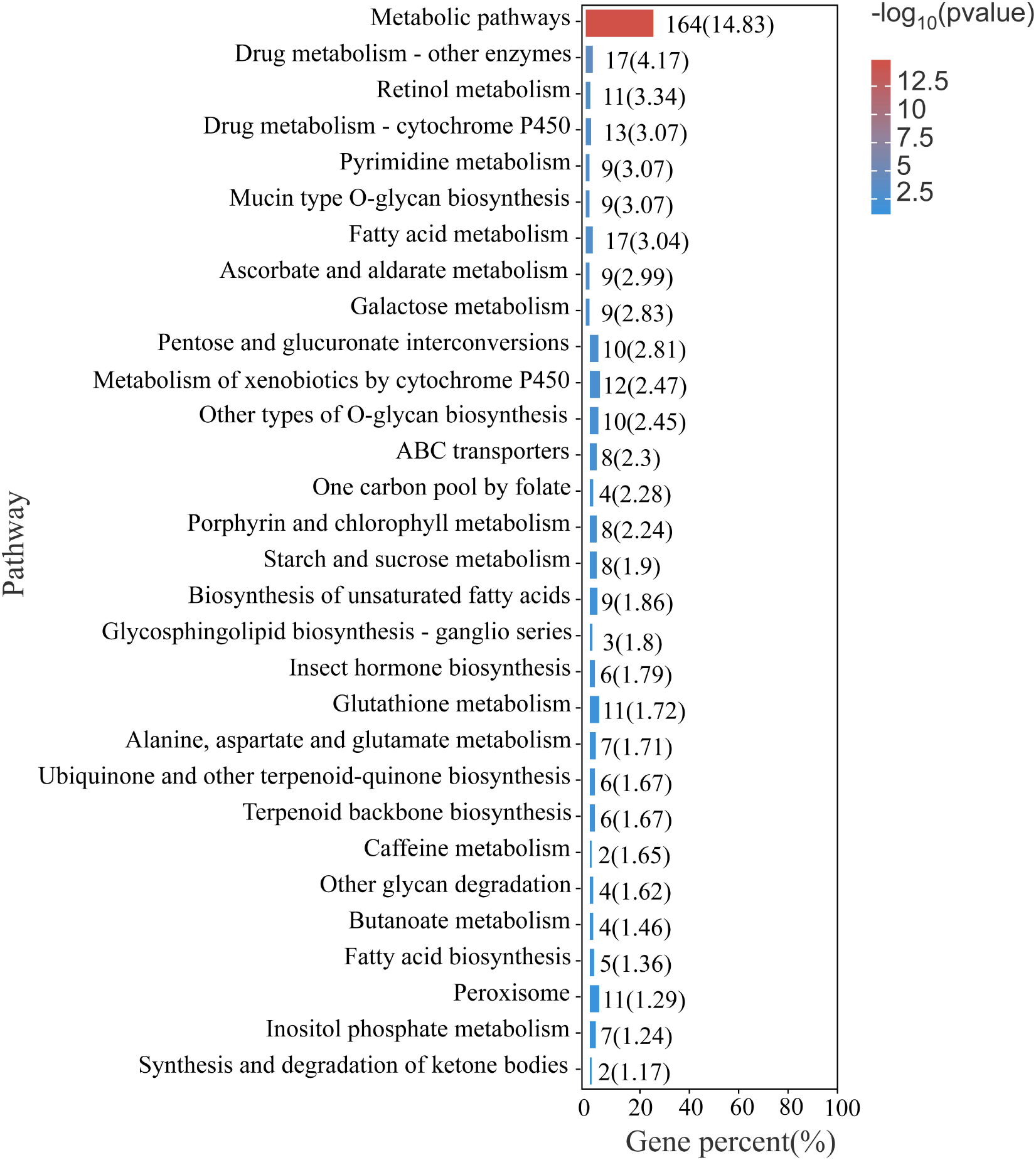
Top 20 of KEGG enrichment analysis of all DEGs.

### 3.5 KEGG enrichment analysis of differentially accumulated metabolites

The analysis indicated that the significantly enriched pathways (q < 0.05) were mainly associated with the amino acid metabolism and synapse, including “Alanine, aspartate and glutamate metabolism”, “Biosynthesis of amino acids”, “Arginine biosynthesis”, “D-Glutamine and D-glutamate metabolism”, “Synaptic vesicle cycle”, “Aminoacyl-tRNA biosynthesis”, “GABAergic synapse” and “Glutamatergic synapse” (Fig 7). Additionally, “Metabolic pathway”, “ABC transporters”, “Microbial metabolism in diverse environments”, “Glyoxylate, dicarboxylate metabolism”, etc. were also significantly enriched. And the differentially accumulated metabolites associated with these pathways include Glutamate, Protoheme, 2,4,5-Trichlorophenol, L-Glutamic acid, 2-Oxoglutarate, and L-Glutamine.

**Fig 7.**
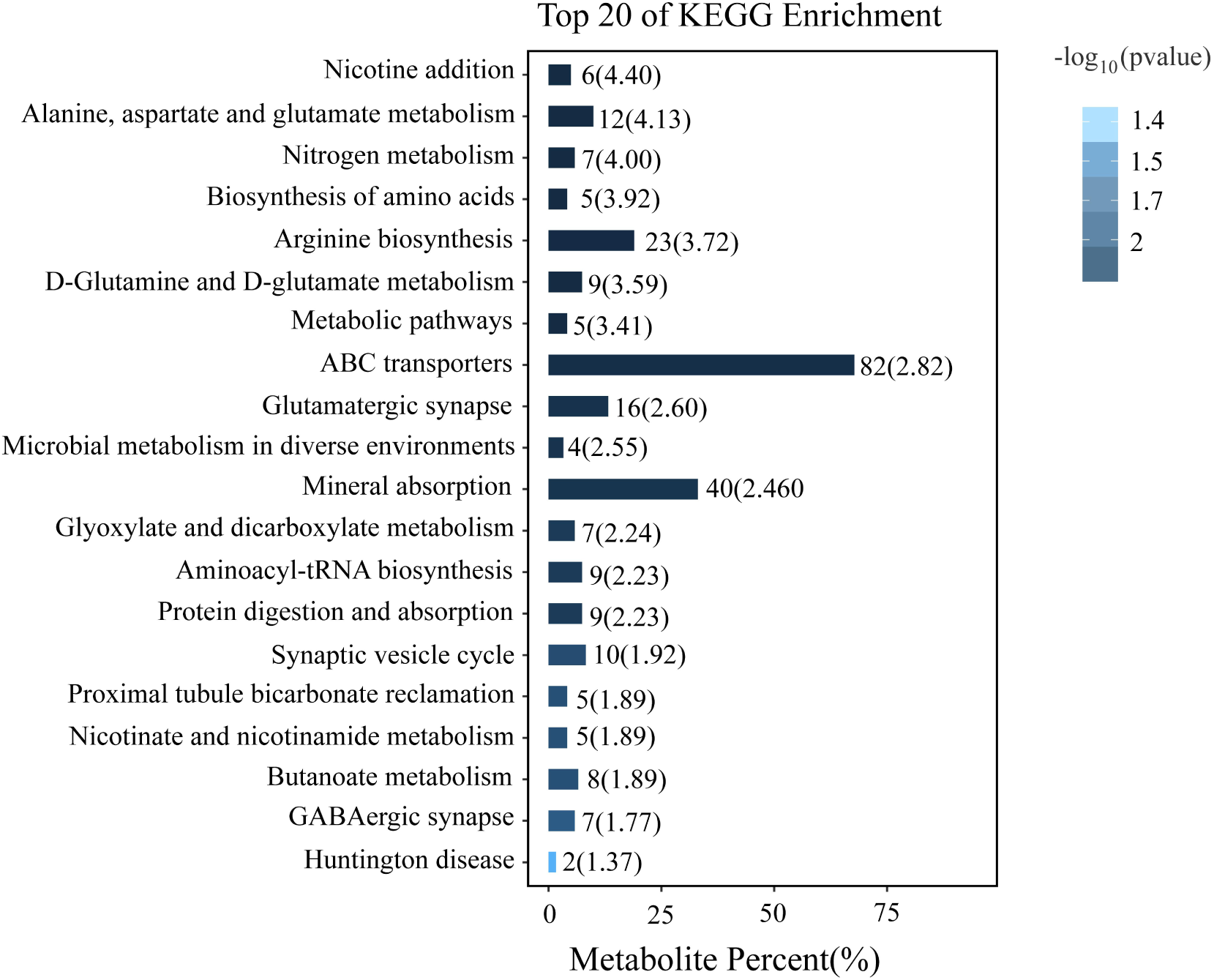
Top 20 of KEGG enrichment analysis of differentially accumulated metabolites.

## 4. Discussion

Hypoxic stress significantly altered Histomorphological features and NKA activity to result in apoptosis (Doonan *et al*., 2008). Meanwhile, our results that the DEGs associated with the transmembrane transport and vesical membrane were significantly enrichment confirmed cell membrane permeability and osmotic pressure are altered under hypoxic stress, which in turn leads to cell membrane shrinkage and cell volume reduction. The apoptotic mechanism in shrimp gills, hepatopancreas, and hemocytes in response to hypoxia has also been studied(Felix-Portillo *et al*., 2016; Nuñez-Hernandez *et al*., 2018; Wang *et al*., 2022). Elevated H_2_O_2_ and ethanol levels, as well as an imbalance in the levels of ROS under hypoxic stress, may result in cell apoptosis in *P. akamusi* (Asai *et al*., 2003; Chen *et al*., 2015; Chen *et al*., 2012; Hasnain *et al*., 1999; Slomiany *et al*., 1997; Souza *et al*., 2003; Young *et al*., 2003). In the present study, the activity of NKA was promoted and showed a fluctuating upward trend in hypoxic stress, but the activity of NKA in the previous study was significantly inhibited and found a significant downward trend in crab and clam (Lucu, 2020; Wang *et al*., 2021). The reason for the different study results may be that acute hypoxia significantly inhibited O_2_^-^ production, which may put the ROS level in an “optimal redox potential range” and thus promote the activity of NKA (Chkadua *et al*., 2022). Furthermore, cellular ion balance was altered by promoting NKA activity, which in turn altered tissue cell permeability, increased cytoplasmic density, and led to cellular tightening (Pratscher *et al*., 2008). The apoptotic vesicles formed were eventually phagocytosed by specialized phagocytes, which may be the reason that the NKA activity decreased significantly under hypoxic stress for 72h. These results, together with the results of the transcriptome, metabolome, biomarkers, and Histomorphological features confirmed that hypoxia-induced apoptosis, which may be an important factor in the ability of *P. akamusi* to tolerate severe hypoxic conditions for long periods. To evaluate the effects of hypoxia on energy metabolism in *P. akamusi*, various parameters were measured. Hypoxic conditions resulted in a significant reduction in glycogen concentrations, indicating a decrease in energy stores (Bao *et al*., 2020). Additionally, metabolites and DEGs associated with lipids, carbohydrate, trehalose, and Glyoxylate cycle were also significantly enrichment, which suggested that they also played an important role in energy metabolic processes under hypoxic stress(Ryabova et al., 2020). The rate of glycogen degradation was slow in acute hypoxic conditions and the longer-term hypoxia causes a further suppression of the energy metabolism in Chironomus larvae, which may be the reason why glycogen no longer decreases after 48h(Choi *et al*., 2001; Hamburger *et al*., 1994; Padilla-Guerrero *et al*., 2011; Popov *et al*., 2005). Lower glycogen content might be closely related to the increasing in LDH activities in the hypoxia group because *P. akamusi*, like *Chaoborus crystallinus* and other macroinvertebrates, might also enhance glycogen decomposition to cope with hypoxia (Englisch *et al*., 1982; Jing *et al*., 2023). In the previous study, ethanol was thought to be the sole end-product of glycogen degradation in *Chironomus riparius* larvae under hypoxia stress (Redecker et al., 1988; Zebe, 1991). The hereby study not only monitored ethanol but also found that ADH activity significantly increased in the hypoxia group. However, this evidence that differentially accumulated metabolites of the microbial metabolism pathway, including 2,4,5-Trichlorophenol and L-Glutamic acid, was significant enrichment and animal cells did not have pyruvate decarboxylase indicated that ethanol was derived from the anaerobic metabolism of the symbiodinium, not from *P. akamusi* larvae (Cook et al., 2010). The use of these enzymes as biomarkers for the quick assessment of stress caused by hypoxia in Chironomidae is supported by the considerably positive connection between the duration of hypoxia and ADH and LDH activity. Our findings provide a new perspective on the aetology of these species by proving that hypoxia is a stressful situation that causes the organism to go into a compensatory response.

The main contribution of acute hypoxia stress to *P. akamusi* may be oxidative stress. The hereby study revealed that extreme hypoxia (DO 0.39 mg O_2_ L^−1^) significantly inhibited the generation of O^−^_2_ and thereby the limited activity of the SOD (Giordano, 2005). According to prior research, mild hypoxia (DO 4 mg O_2_ L^−1^) increased SOD activity while severe hypoxia (DO 2 mg O2 L) had no discernible impact on SOD (Suzuki *et al*., 2019). However, SOD activity was suppressed in an anoxic environment for 8 hours, which is consistent with the findings of our experiment (de Oliveira *et al*., 2005). In essence, the DO concentration and duration of the hypoxia affected the level of SOD. (Lavarías *et al*., 2017). In contrast to SOD, the oxidative stress biomarkers CAT was significantly promoted in the acute hypoxia exposure, and similar results were observed in crab and shrimp(García-Triana *et al*., 2010; Hou *et al*., 2021; Trasviña-Arenas *et al*., 2013). The reason for these results may be that lactate produced by *P. akamusi* can be used as a substrate to generate H_2_O_2_ catalyzed by L-α-hydroxy acid oxidases and ethanol, produced by symbiodinium, also stimulate the function of peroxidase in CAT (Giordano, 2005). Our results that DEGs associated with oxidoreductase and drug metabolism were significantly enrichment also verified the exploration. In addition, significant enrichment of cytochrome P450 in DEGs, including (N-oxide forming)/ hypotaurine monooxygenase (FMO) and glucuronosyl transferase (UGT), indicated that cytochrome P450 also played an important role in promoting ethanol metabolism (Meskar *et al*., 2001). In the actual freshwater ecosystem, dissolved oxygen (DO) concentration fluctuations due to anthropogenic pressure and Global warming were great on spatial and temporal scales (Baoligao *et al*., 2016; Hladyz *et al*., 2011; Pardo *et al*., 2016; Watts *et al*., 2018). However, the DO concentration was the same in this study, thus future investigations may require different combinations of DO concentration.

## 5. Conclusion

We combined histomorphological features and biomarkers with the transcriptome and metabolome to answer the question of why *P. akamusi* can tolerate acute hypoxia stress for long periods. Our results confirmed the altered cell permeability and apoptosis which was induced by H_2_O_2_ and ethanol, regulated by NKA, DEGs, and differentially accumulated metabolites. Research results indicated that lipids and trehalas, besides glycogen, also play an important role in energy metabolism under hypoxic stress. And there may be a glyoxylate cycle in *P. akamusi* or symbiodinium, which converts fatty acid into carbohydrate to cope with insufficient energy. Ethanol produced by the anaerobic respiration of symbiodinium was catalyzed by ADH, CAT, and cytochrome P450 to weaken its harm to the organism, and its content also reflects the level of hypoxia in the environment. While severe hypoxia inhibited SOD activity, lactate, and ethanol promoted CAT activity. The hereby study provided valuable information to understand how *P. akamusi* responded to hypoxic environments in eutrophic lakes and can also help the development of sensitive biomonitoring methods.

### CRediT authorship contribution statement

Yao Zhang & Qing-Ji Zhang contributed equally to this work. Yao Zhang: Conceptualization; Software; Methodology; Formal analysis; Writing – original draft; Data curation; Validation. Qing-Ji Zhang: Conceptualization; Software; Methodology; Formal analysis; Writing – original draft; Data curation; Validation. Wen-bin Xu: Methodology; Formal analysis. Wei Zou: Resources; Validation. Xian-Ling Xiang: Resources; Validation; Supervision. Zhi-Jun Gong: Conceptualization; Methodology; Resources; Writing – review & editing; Supervision; Funding acquisition. Yong-Jiu Cai: Visualization; Resources; Validation; Project administration; Writing – review & editing; Supervision; Funding acquisition; Project administration.

## Acknowledgments

This work was jointly supported by the National Natural Science Foundation of China (Grant 32071572), the Youth Innovation Promotion Association CAS (2020316), and the Science and Technology Planning Project of NIGLAS (NIGLAS2022GS02).

